# Spatiotemporal Dynamics of *Aedes aegypti* and *Culex quinquefasciatus* populations in Miami-Dade County, Florida

**DOI:** 10.1101/2025.10.01.679884

**Authors:** Chalmers Vasquez, Laura C. Multini, John-Paul Mutebi, Ethan SeRine, Megan D. Hill, Maria Litvinova, Marco Ajelli, André B.B. Wilke

## Abstract

Millions of United States residents live where arbovirus vectors like *Aedes aegypti* and *Culex quinquefasciatus* are abundant, and the risk of local outbreaks is amplified by viruses introduced via infected travelers. This threat is well-established: West Nile virus is already endemic in most of the country, and locally acquired dengue outbreaks are occurring with an increasing frequency. Therefore, identifying temporal trends in mosquito abundance and areas conducive to their proliferation is essential for public health preparedness and response planning. This study aims to characterize the spatiotemporal dynamics of *Ae. aegypti* and *Cx. quinquefasciatus* populations in Miami-Dade County, Florida. We analyzed eight years (2017–2024) of mosquito surveillance data from 308 mosquito traps operating across the county. We characterized the spatiotemporal distribution of female *Ae. aegypti* and *Cx. quinquefasciatus* and identified persistent areas of high mosquito abundance (hotspots) using local spatial analysis. A total of 399,418 *Ae. aegypti* and 1,250,879 *Cx. quinquefasciatus* were collected. The two species showed distinct and opposing seasonal patterns: *Ae. aegypti* abundance peaked during the summer wet season, whereas *Cx. quinquefasciatus* peaked in the winter dry season. Our analysis identified spatially consistent hotspots for both species, with some traps classified as hotspots in over half the years studied. The consistent seasonality of the two species and detection of hotspot areas across years provides operational value for long-term monitoring, evaluation of control interventions, and targeted resource allocation. As arboviruses continue to pose a public health risk in urban environments such as Miami-Dade County, the ability to anticipate and respond to vector population fluctuations is instrumental for effective prevention and control.

## Introduction

Millions of United States residents live in areas where *Aedes aegypti*, the primary vector of dengue, Zika, and chikungunya viruses, and *Culex quinquefasciatus*, the primary vector of West Nile virus (WNV) and St. Louis Encephalitis virus (SLEV), are established [1–3]. The risk of local transmission from these mosquitoes is not theoretical. WNV is now endemic in the United States [4], having been documented in 96% of counties since its introduction in 1999 and having caused an estimated ~7 million infections [4,5]. Similarly, locally acquired dengue infections are increasingly reported, with recent outbreaks in Florida, California, and Arizona, highlighting a growing public health threat [6,7]. Specifically, in Miami-Dade County, Florida, 314 locally transmitted dengue virus infections were reported between 2010 and 2024 [8], although CDC estimates suggest that actual case numbers may be substantially higher due to underreporting – estimated reporting rate between 1.0% and 4.8% [9].

While essential for mitigating arboviral disease threats, mosquito control operations are often reactive, often triggered by disease cases or public complaints [10]. A shift toward a more proactive strategy, where interventions can be planned in advance, must be underpinned by a foundational understanding of long-term vector dynamics. This involves identifying consistent seasonal trends for key vector species and persistent high-risk areas (“hotspots”) that can form the basis for data-driven operational planning.

A key step toward proactive control is to analyze historical mosquito surveillance data to identify consistent spatial and temporal patterns in vector populations, including the identification of areas with substantially higher mosquito abundance. To provide this evidence for Miami-Dade County, Florida, we analyzed eight years of surveillance data (2017–2024). Our objectives are to: (i) characterize the spatiotemporal distribution of *Ae. aegypti* and *Cx. quinquefasciatus* and (ii) identify hotspot areas for their proliferation to proactively inform mosquito surveillance and control efforts.

## Materials and Methods

In this study, we used 277 BG-Sentinel 2 traps (Biogents AG, Regensburg, Germany) and 31 CDC light traps that are part of the Miami-Dade County mosquito surveillance system (Fig. 1). Each trap is deployed once weekly for 24 hours over approximately 50 weeks per year. Traps are placed in shaded areas protected from direct sunlight, wind, and precipitation to optimize mosquito collection. Collected mosquitoes are morphologically identified to species using standard taxonomic keys [11]. Surveillance protocols remained consistent throughout the study period, with traps deployed at fixed locations using the same trap models and CO_2_ as bait. BG-Sentinel and CDC traps baited with CO_2_ primarily attract host-seeking female mosquitoes [12] and are not specifically designed to attract males. Although some male mosquitoes were collected, they were considered accidental catches and were excluded from the analyses. Even though the Miami-Dade Mosquito Control surveillance system collected several mosquito species from 2017 to 2024, this study focused on the two primary vector species of epidemiological importance: *Ae. aegypti* and *Cx. quinquefasciatus*.

**Figure 1.**
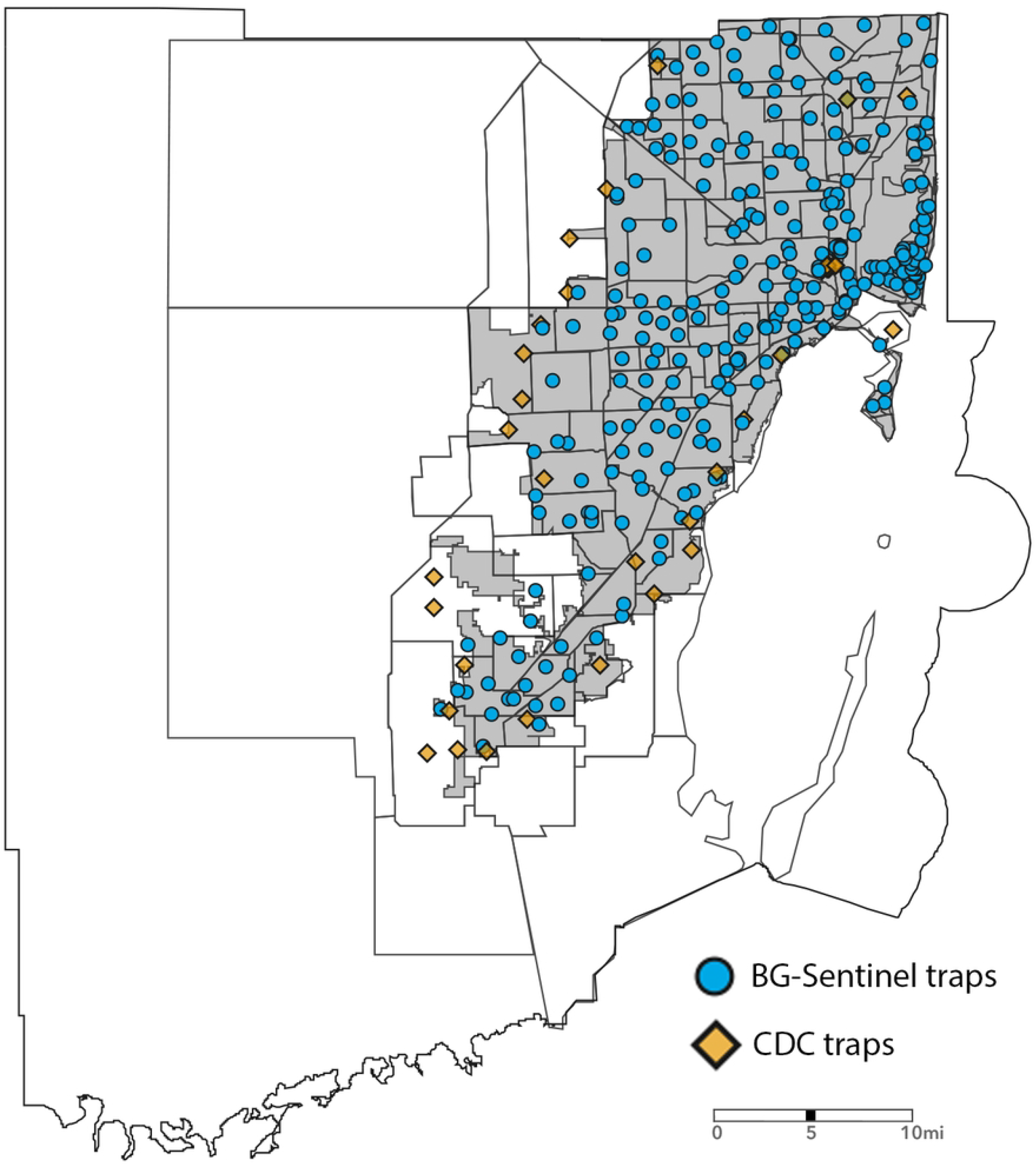
Spatial distribution of traps in the Miami-Dade Mosquito Control Surveillance System, 2024. BG-Sentinel traps are shown as blue circles, and CDC traps as yellow squares.

Monthly species totals were paired with sampling effort, defined as the number of trap-nights per month (calculated by multiplying the number of traps deployed by the number of nights they were operated). Mean female mosquitoes per trap-night were calculated for the entire study period.

The identification of areas conducive to mosquito proliferation serves three main objectives: (1) identify resources at the local scale responsible for supporting mosquito populations; (2) provide an early warning system for increased mosquito activity; and (3) inform targeted control strategies to mitigate mosquito proliferation. Therefore, to identify areas conducive to the proliferation of *Ae. aegypti* and *Cx. quinquefasciatus*, we created a 4-km buffer around each trap and calculated the number of traps within each buffer, along with the total and mean number of mosquitoes collected by each trap in the buffer. This buffer size ensured inclusion of a sufficient number of neighboring traps (mean: 15 per buffer) to enable compelling statistical analysis while maintaining operational spatial resolution. To calculate the mean mosquito abundance per buffer, we removed outliers using a standard method based on 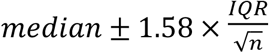 where *IQR* is the interquartile range and *n* is the number of observations in the buffer, as in Wilke et al. [13]. Traps were classified as hotspots if with an average mosquito count above the 97.5th percentile of their respective buffer. All analyses were conducted in R (version 4.2.2).

## Results

From 2017 to 2024, a total of 399,418 *Ae. aegypti* and 1,250,879 *Cx. quinquefasciatus* mosquitoes were collected in Miami-Dade County, Florida (Table 1). Annual counts and mean abundance per trap-night showed consistent patterns, with *Cx. quinquefasciatus* remaining more abundant than *Ae. aegypti* throughout the study period. Yearly relative abundance of *Ae. aegypti* ranged from 35,612 to 64,570 mosquitoes, while *Cx. quinquefasciatus* ranged from 96,154 to 203,223 mosquitoes. Mean values for *Ae. aegypti* ranged from 3.39 to 6.06 mosquitoes per trap-night, while *Cx. quinquefasciatus* ranged from 9.92 to 21.33. The overall mean abundance was 4.25 mosquitoes per trap-night for *Ae. aegypti* and 13.46 for *Cx. quinquefasciatus*.

**Table 1.**
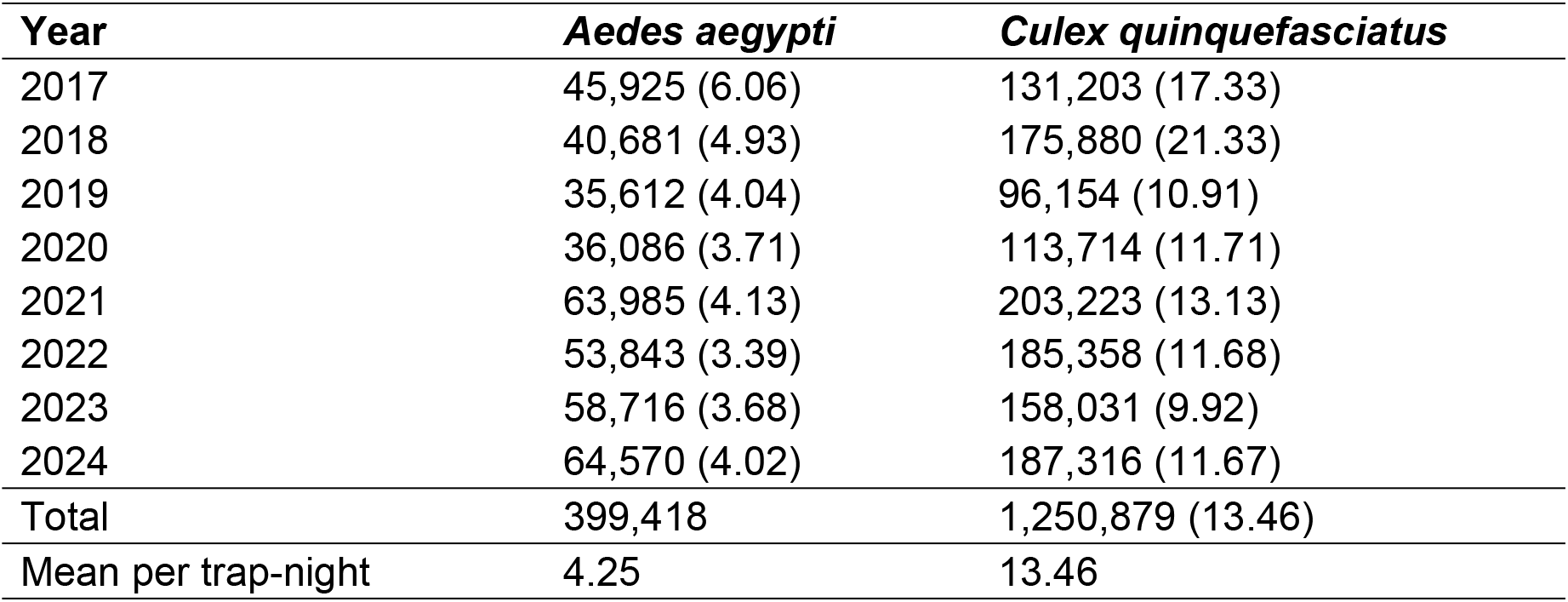
Relative abundance of *Aedes aegypti* and *Culex quinquefasciatus*. Mean number of mosquitoes per trap-night shown in parentheses.

The seasonal population dynamics of *Ae. aegypti* and *Cx. quinquefasciatus* showed consistent temporal variations across the study period (Fig. 2). *Aedes aegypti* showed a seasonal pattern characterized by higher abundance during the summer months (wet season), whereas *Cx. quinquefasciatus* reached peak abundance levels in the winter months (dry season). Throughout the years, *Cx. quinquefasciatus* remained more abundant overall compared to *Ae. aegypti*. Nonetheless, both species were consistently present and collected in relatively high numbers throughout the year, suggesting continuous breeding and survival across seasons. These findings highlight species-specific differences in seasonal dynamics and underscore the importance of year-round surveillance to effectively monitor mosquito vector populations.

**Figure 2.**
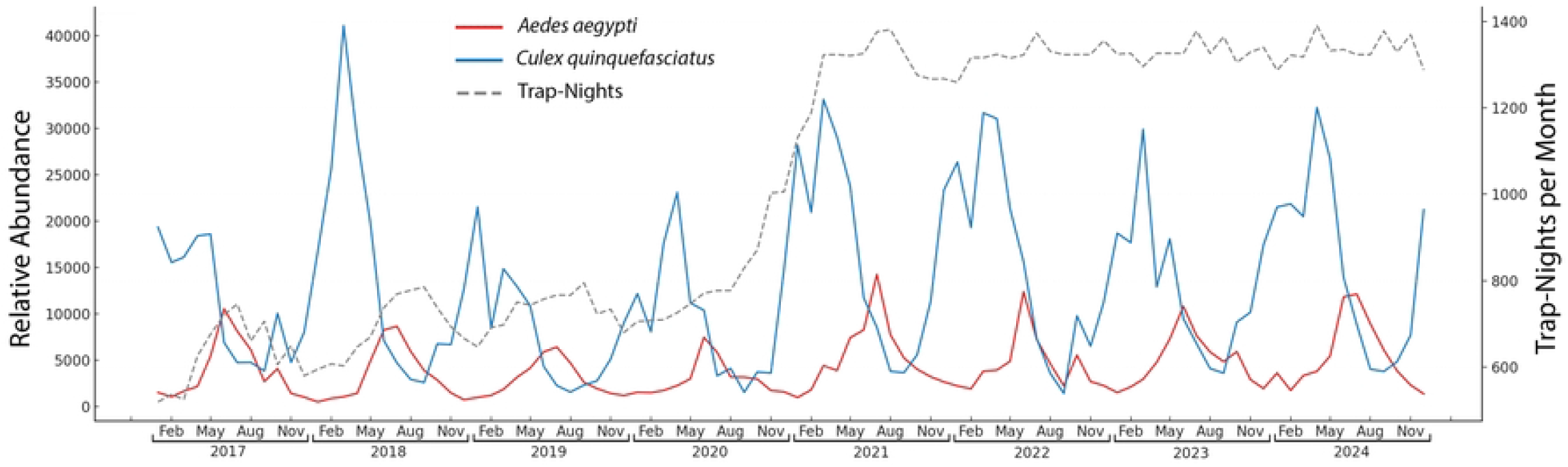
Monthly number of *Aedes aegypti* (red) and *Culex quinquefasciatus* (blue) collected in Miami-Dade County, Florida. The number of trap-nights per month is indicated by the dashed line.

Among the 308 traps included in this eight-year study, an average of 36 traps were classified as hotspots for *Ae. aegypti* (Fig. 3). Of these, 3 traps were identified as hotspots in all eight years, 2 traps in seven years, 4 traps in six years, and 7 traps in five years. Similarly, an average of 34 traps was identified as hotspots for *Cx. quinquefasciatus*, with 3 traps classified as hotspots in all eight years, 4 traps in seven years, 7 traps in six years, and 4 traps in five years.

**Figure 3.**
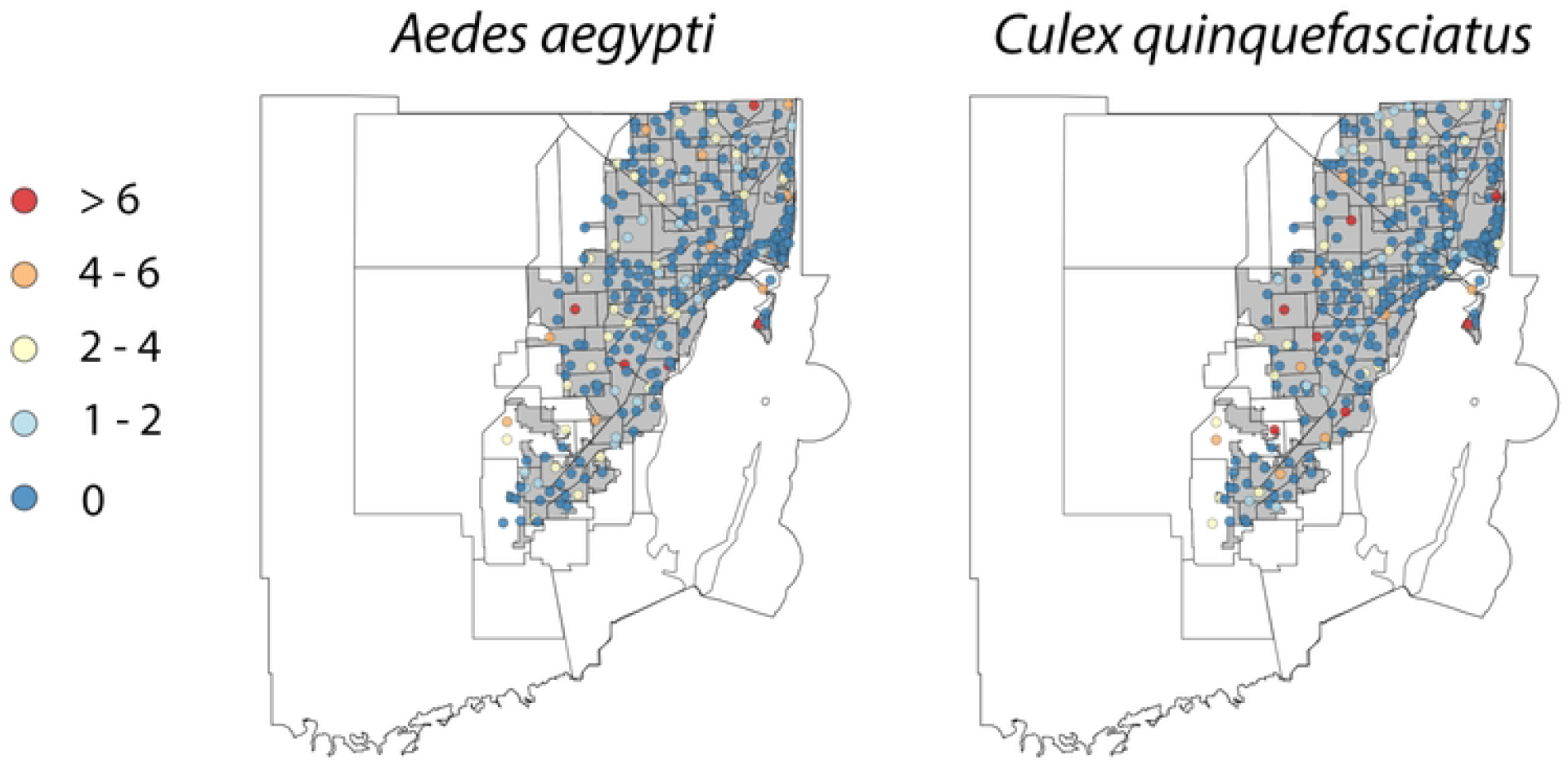
Spatial distribution of hotspots for female *Aedes aegypti* and *Culex quinquefasciatus* in Miami-Dade County, Florida, from 2017 to 2024, based on relative mosquito abundance compared to other traps within a 4 km buffer radius. Colors represent the number of times a trap has been classified as a hotspot during the study period.

## Discussion

The characterization of the spatiotemporal distribution of *Ae. aegypti* and *Cx. quinquefasciatus* in Miami-Dade County, Florida, is essential for identifying hotspot areas for mosquito proliferation to guide control operations and improve arbovirus outbreak preparedness and response. Our results show that both *Ae. aegypti* and *Cx. quinquefasciatus* have well-established seasonal patterns, with *Ae. aegypti* peaking in the summer months and *Cx. quinquefasciatus* in the winter months. The underlying drivers of these seasonal dynamics need further investigation, particularly given the relatively stable temperature in Miami-Dade County throughout the year and the lack of a clear-cut alignment of the relative abundance of the two focus species and rainfall, for example, *Ae. aegypti* population consistently peaked in July despite temperature and rainfall being virtually the same in July and August [14].

Seasonal trends of mosquito vector species have been reported elsewhere in the United States. In Maricopa County, Arizona, *Cx. quinquefasciatus* had bimodal peaks aligned with the spring and early fall rainy seasons, while *Ae. aegypti* populations increased during wetter months of the year and declined sharply at the onset of the dry season [13]. In New Orleans, Louisiana, *Ae. aegypti* showed consistent peaks from May to August, whereas *Cx. quinquefasciatus* maintained high abundance year-round, with broad summer peaks [15]. In Los Angeles, California, *Cx. quinquefasciatus* abundance peaked early in July and declined steadily thereafter, while *Ae. aegypti* increased gradually from June, reaching a peak in October before decreasing in November [16].

These results indicate that while both species respond to seasonal environmental changes, the timing, intensity, and persistence of peaks vary by location, reflecting differences in local climatic patterns, habitat availability, and species-specific ecological adaptations. The patterns found in Miami-Dade County of *Ae. aegypti* summer peak abundance and *Cx. quinquefasciatus* winter peak abundance contrasts with patterns in other regions, underscoring the importance of localized, species-specific surveillance to guide targeted vector control efforts.

Our spatial analysis showed consistent spatial clustering of traps with high mosquito abundance. These spatial trends suggest that local environmental and anthropogenic factors influence species distribution and abundance. The temporal persistence of specific hotspots suggests the presence of stable ecological or structural conditions, as well as resource availability, which support sustained mosquito production. These findings further suggest the presence of priority areas for targeted vector control and support the use of persistent hotspots as sentinel sites for early warning and surveillance.

Miami-Dade County remains particularly susceptible to arbovirus transmission due to its climate, urban landscape, and international connectivity [2,17]. Modeling estimates suggest that, assuming a basic reproduction number of 1.5, the introduction of a single asymptomatic infected individual could lead to a 10% probability of an outbreak resulting in at least 40 symptomatic cases, with a median outbreak size of 73 symptomatic infections [18]. These projections underscore the importance of proactive surveillance and targeted interventions in Miami-Dade County.

Historical spatiotemporal entomological data provide a foundation for the development and implementation of proactive mosquito control measures. However, mosquito populations are highly sensitive to short-term fluctuations in temperature and rainfall, complicating efforts to anticipate and mitigate periods of increased abundance and elevated arbovirus transmission risk [19,20]. This variability presents a challenge for public health agencies responsible for implementing timely and effective mosquito control interventions. Current strategies are primarily reactive, typically triggered by confirmed disease cases, increased mosquito trap counts, or public complaints [21,22]. Integrating high-resolution historical entomological surveillance data into routine operations could enhance situational awareness and support a more proactive, data-driven approach [23,24]. This integration could facilitate the early identification of high-risk periods and locations, enabling public health authorities to optimize intervention timing and targeting, improve resource allocation, and increase the overall effectiveness of mosquito control programs.

## Conclusion

Our findings show that the incorporation of high-resolution entomological surveillance data across spatial and temporal scales enables the identification of key areas and periods of elevated mosquito vector abundance. The consistent detection of hotspot areas across multiple years provides operational value by identifying reliable locations for monitoring mosquito population dynamics, assessing the impact of control interventions, and prioritizing resource allocation. As arboviruses remain a persistent public health threat in urban environments such as Miami-Dade County, the capacity to anticipate and respond to mosquito vector population fluctuations is critical for effective outbreak prevention and control. This study presents data that is instrumental for the improvement of situational awareness, guiding targeted vector control operations, and ultimately reducing the risk of local arbovirus transmission.

